# Cas9-directed long-read sequencing to resolve optical genome mapping findings in leukemia diagnostics

**DOI:** 10.1101/2023.07.21.549991

**Authors:** Eddy N. de Boer, Vincent Vroom, Arjen J. Scheper, Lennart F. Johansson, Laura Bosscher, Nettie Rietema, Sabrina Z. Commandeur, Nine V.A.M. Knoers, Birgit Sikkema-Raddatz, Eva van den Berg, Cleo C. van Diemen

## Abstract

Leukemias are genetically heterogeneous, and knowing the type of genetic aberration is clinically important for progression prognosis and choice of treatment. The standard-of-care (SOC) methods in leukemia diagnostics are karyotyping, SNP-array and FISH. Optical genome mapping (OGM) has emerged as their replacement as it can detect different types of structural aberrations simultaneously. However, although OGM can additionally detect much smaller aberrations (500 bp vs. 5–10 Mb with karyotyping), its resolution is too low to define the location of aberrations between two labels and the breakpoints are disputable when labels are not distinct enough. Here, we test whether Cas9-directed long-read sequencing (LRS) can fill this gap.

From an internal Bionano implementation study we selected ten OGM calls either located in low resolution areas or with disputable breakpoints that could not be validated with SOC methods. Per variant we designed crRNAs for Cas9 enrichment, prepared libraries and sequenced them on a MinION / GridION device.

We could confirm all the OGM aberrations with Cas9-directed LRS and the actual breakpoints of the OGM calls were located between 0.2–5.5 kb of the OGM-estimated breakpoints, confirming the high reliability of OGM. Furthermore, we show examples of redefinition of aberrations between labels that enable judgment of clinical relevance.

We show that Cas9-directed LRS can fill the gap of low resolution OGM areas thereby improving the prediction of clinical significance. We envisage that this technique can be an relevant secondary technique in diagnostic workflows including OGM.

## Introduction

Leukemias are clinically and genetically heterogeneous cancers of the white blood cells that originate in the bone marrow. They are caused by somatic genetic aberrations, including single nucleotide variants (SNVs), small indels, aneuploidies, copy number variations (CNVs), inversions, insertions and (complex) chromosomal rearrangements (1,2,3). Classification of the different types of hematologic malignancies in leukemia is based on the morphology of the cells, immunophenotypic profiles and the identification of specific (cyto)genetic abnormalities (4). As the type of genetic aberration(s) a patient carries in their bone marrow is one of the factors that determine progression prognosis and choice of treatment (5) a rapid and comprehensive genetic diagnosis is critical.

Optical genome mapping (OGM) (Bionano Genomics, San Diego, CA) is an emerging technique that has the potential to detect structural aberrations in different types of leukemia (6). OGM images long (on average 300 kb), directly labeled fragments of genomic DNA. Based on their labels, these images are either directly aligned to a reference genome or *de novo*-assembled before alignment. Several studies using OGM have confirmed that it can identify a wide variety of previously detected aberrations in unselected cohorts of patients with various types of hematologic malignancies (e.g.,7,8).

Although OGM can detect aberrations up to 500 bp with *de novo-*assembly (Bionano, 30110 Rev K), the resolution may be too low as OGM is not able to define the exact location of deletions and duplications that fall between two labels and OGM breakpoints are disputable when the label pattern is not distinctive enough. The exact characterization of aberrations can, however, have significant clinical relevance, for example if an OGM-deleted region between two labels contains a relevant gene and lack of resolution makes it uncertain whether the gene is affected. The current standard-of-care (SOC) methods to validate OGM findings also have limitations. Karyotyping has a limited resolution of 5–10Mb and requires culturing whereas SNP-array has a resolution of 150 kb and can only detect unbalanced aberrations.

In this proof-of-principle study, we tested whether Cas9-directed long-read sequencing (LRS) can be used to characterize aberrations identified with OGM when single-basepair-level resolution is required and/or the resolution of SOC methods is insufficient to confirm OGM detected aberrations.

## METHODS

### Sample selection and workflow

For an internal OGM implementation study, we prospectively collected 18 bone marrow aspirates (BMA) taken from patients with different reasons for referral, a collection set up with a maximum chance of detection of diverse types of aberrations. The inclusion criterion was sufficient BMA to perform OGM alongside current diagnostics. OGM and follow-up experiments were performed in all samples in accordance with the regulations and ethical guidelines of the UMCG.

OGM aberrations were detected as described in Table 1 and Supplemental Information 3. The OGM identified aberrations were then compared to the variants identified using the SOC diagnostic methods. When possible, additional OGM findings were confirmed with the SOC methods karyotyping and the Infinium Global Screening Array-24 v3.0-EA-MD (SNP-Array) (Illumina, San Diego, CA), according to standard protocols. For this proof-of-principle study, we selected samples for Cas9-directed LRS (Oxford Nanopore Technologies, Oxford, UK) when we needed to redefine the breakpoints to improve characterization of the aberration or confirm additional findings that could not be detected with the SOC methods (Table 1).

**Table 1.** Selection of OGM findings for Cas9-directed long-read sequencing.

| ID | Referral reason | OGM (splitted per aberration) | SOC | Identified with SOC | Detection problem |
| --- | --- | --- | --- | --- | --- |
| BMA1 | AML/MDS/MPN | ogm[GRCh37]1q21.1q21.1(144452084_249237532)x2~3 | Karyotyping | yes |  |
| BMA2 | CML | ogm[GRCh37]t(9;22)(q34;q11.2)(133619362_23603312) | Karyotyping | yes |  |
| BMA8 | CML | ogm[GRCh37]t(9;22)(q34;q11.2) | Karyotyping | yes |  |
| BMA16 | HCL (CLL) | ogm[GRCh37](X)x1,(1-22)x2 | Karyotyping | yes |  |
| BMA16 | HCL (CLL) | ogm[GRCh37]fus(8;8)(p22;q13.2) | Karyotyping (1:250 metaphases) | yes |  |
| BMA15 | MDS/AML | ogm[GRCh37] fus(5;5)(q14.2;q33.3) | Karyotyping, SNP-arrays | yes |  |
| BMA8 | CML | ogm[GRCh]22q11.23;q12.1(23,734,214;29,424,129)x1 | SNP-arrays | yes |  |
| BMA9 | MDS | ogm[GRCh37]chr21x3 | SNP-arrays | yes |  |
| BMA10 | MDS/MPN | ogm[GRCh37]9p24.3p13.3(1_35889394)x2 hmz | SNP-arrays | yes |  |
| BMA10 | MDS/MPN | ogm[GRCh37]14q11.2q12(20421719_32361050)x1 | SNP-arrays | yes |  |
| BMA12 | (B)-CLL | ogm[GRCh37]chr12x3 | SNP-arrays | yes |  |
| BMA13 | MDS | ogm[GRCh37] 20q11.22q13.13(34259243_49502193)x1 | SNP-arrays | yes |  |
| BMA16 | HCL (CLL) | ogm[GRCh37]8p23.3p22(11805_15540773)x1~2 | SNP-arrays | yes |  |
| BMA1 | AML/MDS/MPN | ogm[GRCh37]17p13.1(7,545,377_7,588,071)x1 | Could not be confirmed | no | aberration located in between 2 OGM labels |
| BMA3 | AML/MDS | ogm[GRCh37]17p13.1(7,545,377_7,588,071)x1 | Could not be confirmed | no | aberration located in between 2 OGM labels |
| BMA6 | MDS | ogm[GRCh37]17p13.1(7,545,377_7,588,071)x1 | Could not be confirmed | no | aberration located in between 2 OGM labels |
| BMA17 | MDS | OGM[GRCh37]ins(12p13.2)(11,889,007_11,895,594) | Could not be confirmed | no | aberration located in between 2 OGM labels |
| BMA16 | HCL (CLL) | ogm[GRCh37]t(14;17)(q32.22;q25.3)(106,249,815;80,915,618) | Could not be confirmed | no | disputable breakpoint |
| BMA7 | MDS/MF | ogm[GRCh37]inv(3)(q25.33q26.2)(160,014,744_168,882,939) | Could not be confirmed | no | resolution SOC too low |
| BMA7 | MDS/MF | ogm[GRCh37]3q25.33(159,902,689-160,014,744)x1 | Could not be confirmed | no | resolution SOC too low |
| BMA7 | MDS/MF | ogm[GRCh37]3q26.2(168,882,939-168,907,480)x1 | Could not be confirmed | no | resolution SOC too low |
| BMA9 | MDS | ogm[GRCh37]inv(15)(q24.1q24.1)(72,959,741_74,362,190) | Could not be confirmed | no | resolution SOC too low |
| BMA9 | MDS | ogm[GRCh37]3p13(71086423_71375386)x1 | Could not be confirmed | no | resolution SOC too low |

Aberrations detected using optical genome mapping (OGM) in a selected group of 18 bone marrow aspirates (BMA) with varying reasons for referral are depicted in this table. The table indicates if these aberrations were confirmed with the standard-of-care (SOC) methods (yes, no) and, if not, the detection problem is mentioned. Abbreviations referral reason: AML = acute myeloid leukemia, MDS = myelodysplastic syndrome, MPN = myeloproliferative neoplasms, CML = chronic myeloid leukemia, HCL = hairy cell leukemia, CLL = chronic lymphoblastic leukemia, (B)CLL = B-cell acute lymphoblastic leukemia, MF = myelofibrosis.

### DNA extraction, qualification and quantification

The input material for Cas9-directed LRS was the remaining Bionano-isolated ultra-high molecular weight DNA or Maxwell RSC (AS1400) (Promega, Madison, WI) or Nanobind CBB (Pacific Biosciences, Menlo Park, CA) extracted DNA. DNA length, quantity and purity were checked with FemtoPulse pulsed-field capillary electrophoresis (>30 kb) (Agilent, Santa Clara, CA), Nanodrop spectrophotometry (OD 260/280 ∼1.8, OD 260/230 2.0-2.2) (ThermoFisher Scientific, Waltham, MA) and Qubit (ThermoFisher).

### crRNA design for Cas9 enrichment

We used the CHOPCHOP web browser (9,10,11) to search for candidate crRNA sequences. The ONT instructions (targeted, amplification-free DNA sequencing using Crispr/CAS, version ECI_S1014_v1_revE_11Dec2018) were used in combination with Genome build GRCh37. Efficiency scores of crRNAs were calculated according to “Doench et al. 2014 -only for NGG PAM” (12). The downloaded results were filtered as recommended by ONT: GC: 40-80%, self-complementarity score 0, retain efficiency score >0.3 (Cas9 only), retain candidates with mismatches MM0=0, MM1=0, MM2=0, MM3 least possible. The crRNAs we designed are listed in Supplemental Information 4 and were ordered form IDT (IDT, Coralville, IA (IDT) as Alt-R^TM^ Cas9 crRNAs.

### Enrichment, sample-preparation and sequencing

We used the ligation sequencing Cas9 enrichment protocol (SQK-CS9109, ONT) to enrich the regions of interest and prepare the samples for sequencing, following the manufacturer’s instructions. The protocol was started with an input of 1–5 µg HMW DNA. The library was purified with AMPure XP beads (A63881) (Beckman Coulter^TM^, Fullerton, CA). An R9 MinION flowcell (FLO-MIN114) was primed, loaded and run on a MinION / GridION device, following ONT instructions.

### Cas9 data analysis

Basecalling was performed using MinKNOW v22.10.5 (ONT). Fastq files passing quality metrics were merged into a single fastq file and aligned with minimap2 v2.24 (13). Sniffles2 v2.0.7 (14) was used for SV calling. In addition, bam-files were viewed within IGV-viewer v2.14.1 (15). In association with a tumor cytogeneticist, we assessed the potential pathogenicity of the variants based on the literature and the WHO classification of tumors of hematopoietic and lymphoid tissues (4,16).

## RESULTS

### Selection of samples for Cas9-directed long-read sequencing

The implementation study resulted in detection of 23 aberrations with OGM. Of these, 13 were confirmed with the results of the SOC methods, some performed as part of the diagnostic process. For these variants, it was not necessary to know the exact location of the breakpoints, and no Cas9-directed LRS confirmation was performed. This left 10 aberrations that could not be confirmed using the SOC methods and were selected to follow up with Cas9-directed LRS (Table 1).

### Confirmation of additional OGM findings and redefinition of the breakpoints with Cas9-directed long-read sequencing

We could detect each selected aberration using Cas9-directed LRS. We used at the maximum one R9 MinION flowcell (FLO-MIN114) per aberration, resulting in 1-25 LRS reads covering the aberration.

#### Redefinition of the breakpoints

In three patients (BMA1, BMA3 and BMA6), OGM detected a heterozygous 1.2 kb deletion in a 42 kb region located between two OGM labels (ogm[GRCh37]17p13.1(7,545,377_7,588,071)x1) that includes a part of *TP53* (chr17:7,571,720-7,590,868). Cas9-directed LRS redefined the deletion to a 1.6 kb (chr17:7,564,612-7,566,197) deletion localized adjacent to *TP53* (Figure 1). Because *TP53* is not included in the deletion, this aberration is not considered clinically relevant, and it is also a known CNV in the general population (17).

**Figure 1.**
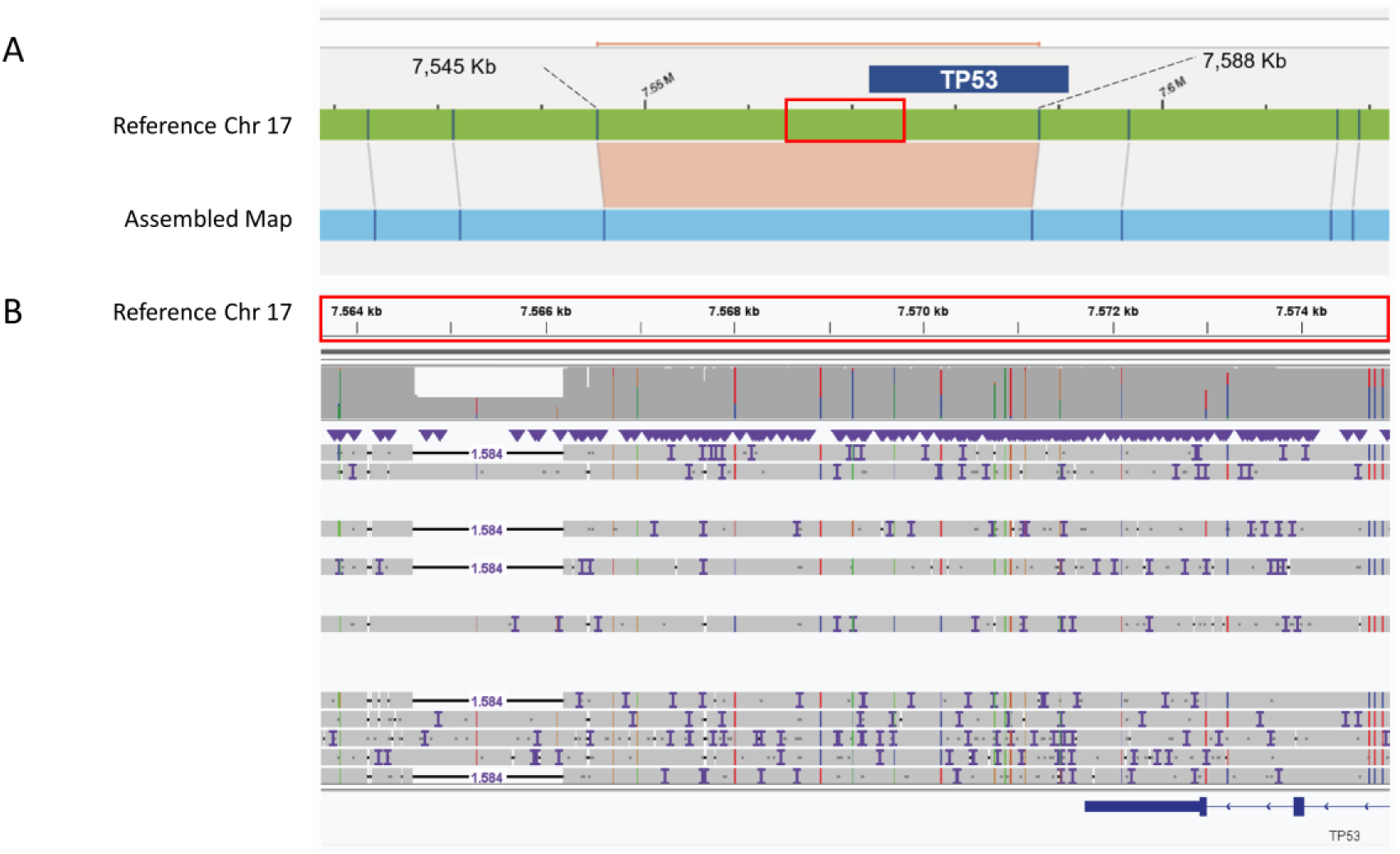
Representative example of an OGM aberration located between two OGM labels. A) In three patients, OGM detected a heterozygous 1.2 kb deletion in a 42 kb region (17p13.1) between two OGM labels that overlaps with *TP53*. B) Cas9-directed long-read sequencing redefined the aberration to a 1.6 kb deletion localized upstream of *TP53*. This aberration, a known CNV, is characterized as not clinically relevant.

In patient BMA17, referred for myelodysplastic syndrome, OGM detected a heterozygous 1.4 kb insertion in a 6.6 kb region located between two OGM labels (ogm[GRCh37]ins(12p13.2)(11,889,007_11,895,594)) that includes *ETV6* (chr12:11,802,788-12,048,325). Cas9-directed LRS redefined the insertion to 1.4 kb. This 1.4 kb insertion is 99% similar to region chr12:11,892,052-11,893,440 (https://genome.ucsc.edu/cgi-bin/hgBlat) and is inserted in intron 1 (chr12:11,892,830-11,894,194) of the NM_001987 transcript of *ETV6*. The localization of this inserted region is deep intronic, and thus it probably has no clinical significance.

In patient BMA16, referred for hairy cell leukemia, OGM detected a balanced translocation between chromosomes 14 and 17 (ogm[GRCh37]t(14;17)(q32.22;q25.3)(106,249,815;80,915,618)) that includes *IGHG1*. The breakpoint is disputable but is relevant for prognosis (Figure 2). The translocation could not be confirmed with SNP-array and karyotyping in mature cells, and the resolution of FISH is too low to precisely localize the breakpoints. Cas9-directed LRS redefined the breakpoints to (chr14:106,114,465;chr17:81,009,018) (Figure 2). The breakpoint on chromosome 14 (106,114,465) is located 88 kb 3’ downstream of *IGHG1* (chr14:106,202,680-106,209,408). This translocation has not been described in the context of leukemia.

**Figure 2.**
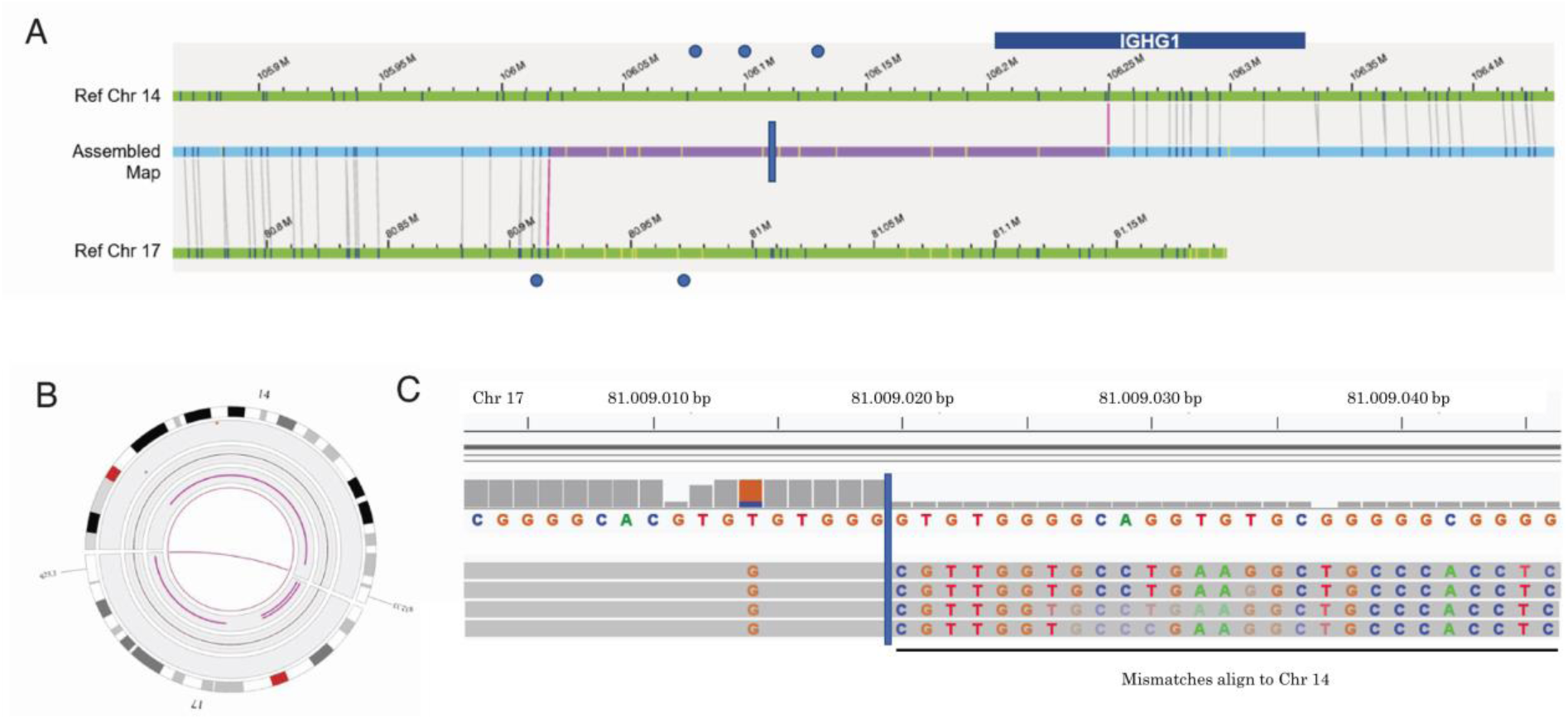
Representative example of an OGM aberration with a disputable breakpoint. OGM detected a translocation between chromosome 14 and 17 (q32.33;q25.3) that includes part of *IGHG1* (A & B). We evaluated the uncertain breakpoint region (purple area, A) and used this information to design crisprs (blue dots, A). Cas9-directed long-read sequencing redefined the breakpoints (blue vertical line, A & C). The chromosome 14 breakpoint is located 88 kb upstream of *IGHG1* (A & C).

#### Increased resolution required for confirmation of an OGM call

In patient BMA7, who was referred for myelodysplastic syndrome/myelofibrosis, OGM detected three aberrations: an 8.9 Mb inversion (ogm[GRCh37]inv(3)(q25.33q26.2)(160,014,744_168,882,939)), a 112 kb deletion of 3q25.33 (ogm[GRCh37]3q25.33(159,902,689-160,014,744)x1) and a 25 kb deletion of 3q26.2 (ogm[GRCh37]3q26.2(168,882,939-168,907,480)x1). Cas9-directed LRS redefined the inversion to 8.9 Mb (160,014,965-168,888,415) and the deletions to 109 kb (159,905,686-160,014,965) and 19 kb (168,888,415-168,907,304), respectively (Figure 3). The redefined breakpoints are located, respectively 0.22, 5.5, 3.0, 0.22, 5.5 and 0.18 kb from the OGM breakpoints. The 19 kb deletion (168,888,415-168,907,304) is part of intron 2 of the NM-004991.3 transcript of *MECOM* and is located 5’ upstream of other transcripts. The 8.9 Mb inversion (160,014,965-168,888,415) includes part of transcript NM-004991.3 (intron 2 till exon 16), which will cause a non-functioning transcript. In the other transcripts, the inversion includes all of *MECOM*. In these transcripts, *MECOM* (q26.2) is repositioned towards the *GATA2* enhancer (3q21) that drives *MECOM* expression (18). However, this is not the previously identified inversion 3 (q21q26) that is associated with leukemia.

**Figure 3.**
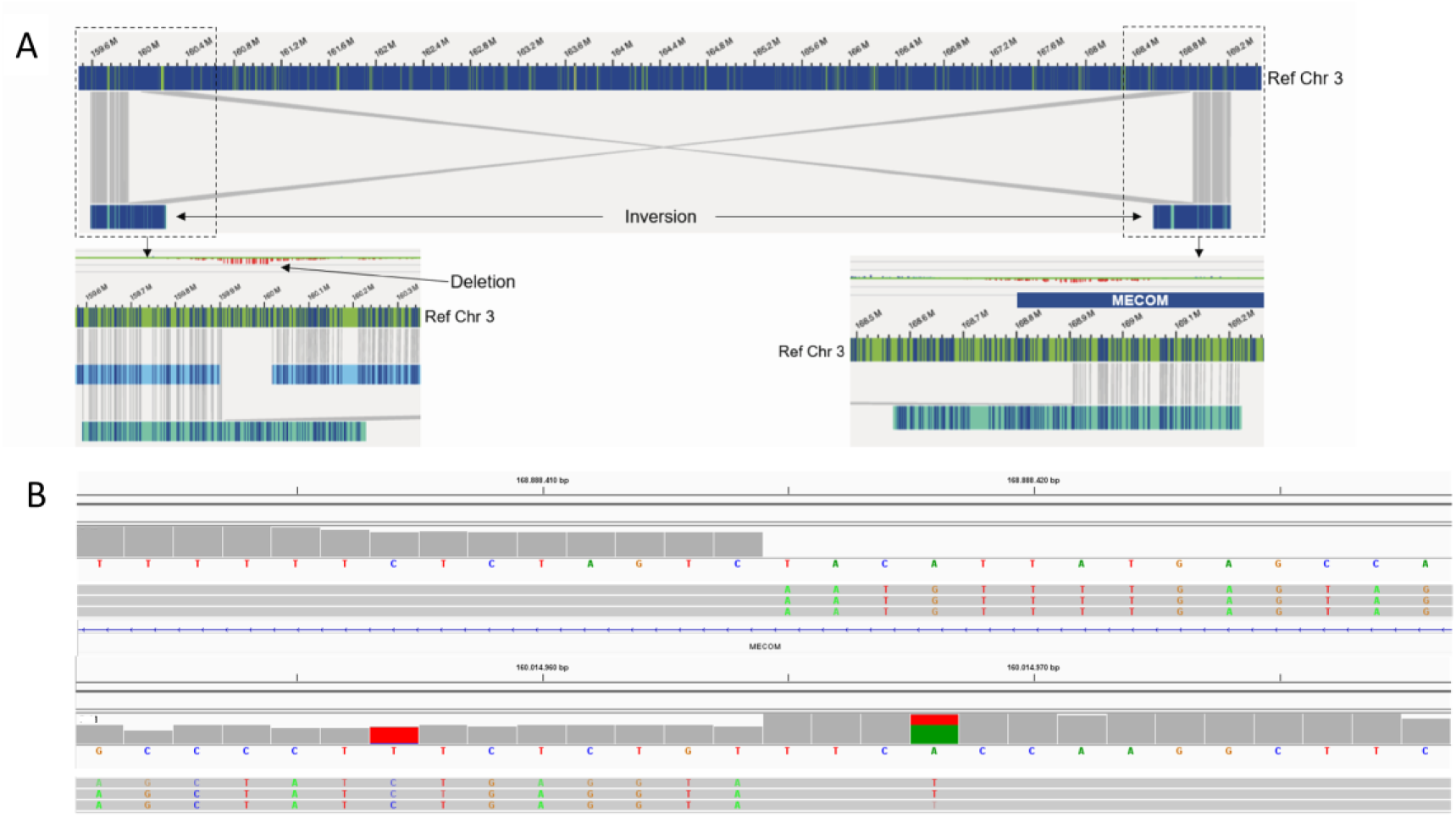
Representative example of finding where SOC resolution is too low. A) OGM detected an 8.9 Mb inversion of chromosome 3(q25.33q26.2) that includes *MECOM*, while two adjacent 112 kb (q25.33) and 25 kb (q26.2) regions were deleted. B) Cas9-directed long-read sequencing redefined the inversion to 8.9 Mb (160,014,965-168,888,415) and the deletions to 109 kb (159,905,686-160,014,965) and 19 kb (168,888,415-168,907,304). The inversion includes part of (NM-004991.3) or all of the *MECOM* transcript.

In patient BMA9, who was referred for myelodysplastic syndrome, OGM detected a heterozygous 289 kb deletion that includes *FOXP1* (ogm[GRCh37]3p13(71,086,423_71,375,386)x1). This aberration, which has an allele frequency of 3%, was only detected with the OGM rare variant pipeline (Supplemental Information 3). Cas9-directed LRS redefined the aberration to a 283 kb (71,087,137-71,370,212) deletion of intron 7 through intron 11 of *FOXP1.* The redefined breakpoints are located 0.7 kb and 5 kb of the OGM breakpoints, respectively. *FOXP1* deletions have been described in acute myeloid leukemia and myelodysplastic syndrome, but the evidence for their clinical relevance is currently not strong (19,20).

Finally, in patient BMA9, OGM detected a 305 kb inserted inversion including the *PML* gene located directly upstream of *GOLGA6B* (ogm[GRCh37]inv(15)(q24.1q24.1)(72,947,038_72,959,738)). In the wild-type situation, *PML* (chr15:74,287,014-74,340,155) is located upstream of *GOLGA6A* (chr15:74,362,198-74,374,891). After closer inspection of the OGM data, the whole 1.4Mb region between *GOLGA6B* and *GOLGA6A* (ogm[GRCh37]inv(15)(q24.1q24.1)(72,959,741_74,362,190)) is inverted, and *PML* is not included in the inversion. The breakpoints are located within or nearby *GOLGA6A* and *GOLGA6B*, which have the same OGM label pattern. *GOLGA6A* and *GOLGA6B* are duplicons with 99.7% similarity. The uniquely mapped Nanopore reads became non-unique at the *GOLGA6B* locus. These reads are also non-uniquely mapping to the *GOLGA6A* locus (recognized by the read-ID).

We identified forward-orientated (mutant) and reverse-orientated (wild-type) adapters at the CRISPR cutting sites. This means that the CRISPR cutting sites were located within the inverted region of the mutant allele, which does confirm the heterozygous inversion. The CRISPRs were designed in the wrong orientation because we initially misjudged the OGM aberration. As a result, we could not redefine the breakpoints. This inversion has not been described in the context of leukemia.

## Discussion

In this proof-of-principle study, we show that Cas9-directed LRS can be used to characterize low resolution areas of OGM at single-basepair-level to improve the interpretation of aberrations. This is exemplified by our redefinition of a heterozygous 1.2 kb deletion in a 42 kb region (17p13.1) that includes *TP53* that fell between two OGM labels. Cas9-directed LRS showed that *TP53* is located upstream of the deletion. Another example was an uncertain OGM breakpoint of a translocation between chromosome 14 and 17 (q32.33; q25.3) where it was uncertain whether *IGHG1* was included. Cas9-directed LRS redefined the breakpoint to 88 kb upstream of *IGHG1.* These two examples demonstrate the relevance of using a high-resolution method like Cas9-directed LRS to follow-up OGM findings to determine the involvement of genes in a structural aberration, and therefore the clinical consequence.

This proof-of-principle study confirms that OGM is very reliable. Other previous studies (e.g. 7,8) did not attempt to confirm OGM findings below the resolution of the SOC methods, whereas we could also confirm such aberrations. Cas9-directed LRS confirmed several different types of OGM aberrations including translocations, deletions, insertions and inversions. In total we detected 23 OGM-aberrations in 18 BMAs. Of these aberrations, we confirmed 5 aberrations with Cas9-directed LRS because the resolution of the SOC methods was too low while we redefined the breakpoints of 5 other OGM-aberrations with insufficient resolution. We show that the actual breakpoints of the OGM calls are located between 0.2–5.5 kb of the OGM-estimated breakpoints, which supports previous reports. Because of this, we expect that standard confirmation of OGM calls with sufficient resolution will not be needed after extensive validation in the laboratories that opt to implement this technology in their workflow. However, in a future diagnostic setting in which the SOC methods will likely be phased out, it will be useful to have Cas9-directed LRS available to confirm potential clinically relevant OGM calls. However, this approach needs extensive further validation, in particular because our study is small and implementation of a new diagnostic workflow needs a prolonged period of validation of variants.

The coverage of the region of interest of Cas9-directed LRS depends on biological variation and technical variables as DNA-quality, crRNA efficiency, sample-prep performance and sequencing performance. Indeed, the number of reads covering the targeted aberration (1-25 reads) was variable and sometimes sub-optimal because of the piloting nature of this study. Still, the coverage of this targeted method is generally higher than a whole genome sequencing approach would yield. In theory, one read with the aberration is sufficient to confirm and redefine the aberration with Cas9-directed LRS. A high coverage would however make it easier to confirm and redefine aberrations, especially when the frequency of somatic cells with the aberration in the sample is low. In this pilot study, we detected a 283 kb *FOXP1* deletion (3% allele frequency) with OGM and confirmed and redefined this aberration with Cas9-directed LRS. If the expected frequency of the aberration is estimated to be relatively high based on the OGM data, it will be possible to decrease the sequencing costs by sequencing more than one sample on one flow-cell. Currently, we observe that we are capturing on average a higher number of reads with the aberration as we are getting better acquainted with Cas9-directed LRS.

The strength of Cas9-directed LRS is that it is an amplification-free method that can access repeat regions, GC-rich areas and homologous regions in contrast to methods requiring PCR, including short-read sequencing and long range PCR (21,22). Additionally, these methods have a limited reach. For example, it would not have been possible to cover the whole 42 kb region between the two OGM-labels in case of the *TP53* region, because it was unknown where the aberration was located. For known and possibly returning yet characterized CNVs long range PCR can be used for confirmation as a possibly cheaper alternative to Cas9-directed LRS. An alternative technique to confirm aberrations is (targeted) short read sequencing, however, it is much more difficult to span the breakpoints with high confidence, in particular with aberrations of low frequency. Therefore, we suggest that Cas9-directed LRS is the most appropriate method for confirming aberrations detected by OGM. A similar approach is, in principle, possible using targeted Pacific Biosciences SMRT sequencing (Menlo Park, CA), but Cas9-directed LRS has the advantage that an amplification-free enrichment is already included in the workflow. As opposed to the PacBio method which needs a separated enrichment.

Any new diagnostic workflow should have a short turnaround time in order to provide the best clinical care in leukemia diagnostics. For this reason, it is important to have readily available validation options for disputed OGM calls. This approach has prognostic value, as we showed for the 42 kb region that included *TP53* that fell between two OGM labels. Because we already know the genomic regions for which OGM has limited resolution, it is possible to design and validate guide-RNAs for known hotspots in advance. The turnaround time of Cas9-directed LRS can then be limited to 1 or 2 working day(s), as is desired for prognosis and treatment in leukemia diagnostics. In a situation where no guide-RNAs are readily available, it will take 1.5–2 weeks to design and order guide-RNAs. However, we expect that guide-RNAs can be stored for years, which will allow extensive libraries to be built up for validation purposes.

We and others have shown that OGM, in combination with Cas9-directed LRS, reveals novel structural abnormalities and can unravel more complex aberrations. At the moment, however, the clinical significance of the aberrations remains unclear. We expect that prediction of clinical significance will improve over time through growing knowledge and enriched variant databases, both of which can be accelerated by cooperation in (inter)national working groups. Although OGM is a method that accesses the whole genome, we are currently forced to limit the region of interest to regions known to be involved in leukemia due to the high number of variants of uncertain significance. In the future, extending of the region of interest may provide additional information for better prognosis and treatment.

In conclusion, OGM in combination with Cas9-directed LRS is a workflow that can characterize (complex) aberrations down to single-basepair-level. This approach will improve the prediction of the clinical significance of OGM-identified aberrations. Although OGM is currently the only method that produces long-reads with the required coverage to resolve somatic complex rearrangements, in some cases, Cas9-directed LRS is needed to fill gaps where OGM resolution is insufficient.

## Funding and Disclosures / Conflict of interest

The authors have no conflicts of interest, including specific financial interests and relationships and affiliations relevant to the subject of the manuscript.

### List of abbreviations

SNV: single nucleotide variants
CNV: copy number variation
OGM: optical genome mapping
SOC: standard of care
LRS: long-read sequencing
BMA: bone marrow aspirates

## Human Genes

TP53

ETV6

IGHG1

MECOM

GATA2

FOXP1

PML

GOLGA6A

GOLGA6B

## Acknowledgements

We thank Kate McIntyre for editing.

## References

1. Grimwade D, Freeman SD. Defining minimal residual disease in acute myeloid leukemia: which platforms are ready for “prime time”? Blood 2014;124(23):3345–3355.

2. Ley TJ, Miller C, Ding L, Raphael BJ, Mungall AJ, Robertson AG et al. Genomic and epigenomic landscapes of adult de novo acute myeloid leukemia. N Engl J Med 2013;368(22):2059–2074.

3. Luthra R, Patel KP, Reddy NG, Haghshenas V, Routbort MJ, Harmon MA et al. Nextgeneration sequencing-based multigene mutational screening for acute myeloid leukemia using MiSeq: applicability for diagnostics and disease monitoring. Haematologica 2014;99:465–73.

4. Khoury JD, Solary E, Abla O, Akkari Y, Alaggio R, Apperley JF et al. The 5^th^ edition of the Worlth Health Organization Classification of Haematolymphoid Tumours: Myeloid and Histiocytic/Dendritic Neoplasms. Leukemia 2022;36(7):1703–1719.

5. Arber DA, Orazi A, Hasserjian R, et al. The 2016 revision to the World Health Organization classification of myeloid neoplasms and acute leukemia. Blood 2016;127(20):2391-2405.

6. Bocklandt S, Hastie A, Cao H. Bionano Genome Mapping: High-throughput, Ultra-Long Molecule Genome Analysis System for Precission Genome Assembly and Haploid-Resolved Structural Variation Discovery. In: Suzuki, Y. (eds) Single Molecule and Single Cell Sequencing. Adv in Exp Med and Biol 2019;1129:97-118.

7. Neveling K, Mantere T, Vermeulen S, Oorsprong M, van Beek R, Kater-Baats E et al. Next-generation cytogenetics: Comprehensive assessment of 52 hematological malignancy genomes by optical genome mapping. Am J Hum Genet. 2021;108(8):1423–1435.

8. Lestringant V, Duployez N, Penther D, Luquet I, Derrieux C, Lutun A et al. Optical genome mapping, a promising alternative to gold standard cytogenetic approaches in a series of acute lymphoblastic leukemias. Genes Chromosomes Cancer 2021;60(10):657–667.

9. Labun K, Montague TG, Krause M, Torres Cleuren YN, Tjeldnes H et al. CHOPCHOP v3: expanding the CRISPR web toolbox beyond genome editing. Nucleic acids research 2019;47(W1):171–174.

10. Labun K, Montague TG, Gagnon JA, Thyme SB, Valen E. CHOPCHOP v2: a web tool for the next generation of CRISPR genome engineering. Nucleic Acids Research 2016;44(W1):272–276.

11. Montague TG, Cruz JM, Cagnon JA, Church GM, Valen E. CHOPCHOP: a CRISPR/CAS9 and TALEN web tool for genome editing. Nucleic acids research 2014;42(W1):401–407.

12. Doench JG, Hartenian E, Graham DB, Tothova Z, Hegde M, Smith I et al. Rational design of highly active sgRNAs for CRISPR-Cas9-mediated gene inactivation. Nat Biotechnol. 2014;32:1262–1267.

13. Li H. Minimap2: pairwise alignment for nucleotide sequences. Bioinformatics 2018;34:3094–3100.

14. Sedlazeck FJ, Rescheneder P, Smolka M, Fang H, Nattestad M, von Haeseler A, Schatz MC. Accurate detection of complex structural variations using single-molecule sequencing. Nat methods. 2018;15(6):461–468.

15. Robinson JT, Thorvaldsdóttir H, Winckler W, Guttman M, Lander ES, Getz G, Mesirov JP. Integrative genomics viewer. Nature Biotechnology 2011;29:24–26.

16. Swerlow SH, Campo E, Harris NL, Jaffe ES, Pileri SA, Stein H, Thiele J: WHO Classification of Tumours of Haematopoetic and Lymphoid Tissues (revised 4^th^ edition). IARC: Lyon 2017.

17. The 1000 Genomes Project Consortium. A global reference for human genetic variation. Nature 2015;526(7571):68-74.

18. Suzuki M, Katayama S, Yamamoto M. Two effect of GATA2 enhancer repositioning by 3q chromosomal rearrangements. IUBMB Life 2019;72(issue1):159–169.

19. L’Abbate A, Lo Cuncolo C, Macri E, Luzzolino P, Mecucci C, Doglioni C et al. FOXP1 and TP63 involvement in the progression of myelodysplastic syndrome with 5q-and additional cytogenetic abnormalities. BMC Cancer 2014;14:396.

20. Klampfl T, Harutyunyan A, Berg T, Gisslinger B, Schalling M, Bagienski K et al. Genome integrity of myeloproliferative neoplasms in chronic phase and during disease progression. Blood 2011;118(1):167–176.

21. Ebbert MTW, Jensen TD, Jansen-West K, Sens JP, Reddy JS, Ridge PG et al. Systematic analysis of dark and camouflaged genes reveals disease-relevant genes hiding in plain sight. Genome Biol 2019;20:97.

22. Hu L, Liang F, Cheng D, Zhang Z, Yu G, Zha J et al. Location of balanced chromosome-translocation breakpoints by long-read sequencing on the Oxford nanopore platform. Front. Genet. 2020; 10:1313.

