## Supplemental Information for "Cas9-directed long-read sequencing to resolve optical genome mapping findings in leukemia diagnostics"

**Supplemental Information 1: the OGM settings.**

Data-analysis for the *de novo* assembly and rare variant analysis (RVA) was optimized by comparing several filter settings (Supplemental Table 1). Settings with a variable number of aberrations are the positive predictive value (PPV) of the intra-fusions, the PPV of the inter-translocations, the presence of the variants in the control group and the masking of CNV-sensitive regions of the genome (Supplemental Table 2 and 3). As hematologic malignancies arise at all ages (Prog Tumor Res. 2016;43:87-100, Juliusson et. al.), we used the least stringent filtering of the aberrations in the control group. Some aberrations were filtered when masking the CNV-enriched regions or selecting the genes of interest using the Access software. For example, an inversion (3q25.33;q26.2) that includes *MECOM* was filtered out due to a bug in the Access v.1.7 software. Further, translocations and fusions without a gene of interest were filtered out even though these aberrations can be of clinical significance. As a consequence, we decided to use settings D (Supplemental Table 1) which had the lowest specificity and highest sensitivity for the final analysis. Inverted aberrations in the genes of interest and disease-causing aberrations without a gene of interest were filtered using a disease-specific BED-file (Supplemental Information 2). For this reason, disease-specific BED files were used to visualize the regions of interest, but not for initial filtering.

The final settings (settings D) are: insertion recommended (confidence 0) and minimum size 500 bp, deletion recommended (confidence 0) and minimum size 500 bp, inversion recommended (confidence 0.7) and minimum size 30 kbp, duplication recommended (confidence -1) and minimum size 30 kbp, intra-fusion recommended (confidence 0.05), and; inter-translocation recommended (confidence 0.05). Detected events were subsequently filtered (less than or equal to 1%) using a database of approximately 300 individuals from 33 different ethnicities with no known genetic disorder available in Access v.1.7. Settings applied to all SV types were: SV-masking = all structural variants, VAF filter min = 0, VAF filter max = 1, SV self-molecule check = SV found in self-molecules, self-molecule count = 5, SV overlapping genes filter = all SVs. CNV filters were: CNV type = all, copy CNV confidence = recommended (0.99), CNV minimum size = 500 kbp, CNV masking filter = all. Aneuploidy filters were: aneuploidy type = all, aneuploidy confidence = recommended (0.95). AOH/LOH filters (*de-novo*) were: AOH/LOH minimum size = 0bp. GRCh37 hg19 (UCSC, Santa Cruz USA (UCSC)) was used as a reference genome.

**Supplemental Table 1: Filter settings tested.**

| | Filterset A | Filterset B | Filterset C | Filterset D | **Filterset E** |
|---|---|---|---|---|---|
| Insertion | 0 | 0 | 0 | 0 | 0 |
| Deletion | 0 | 0 | 0 | 0 | 0 |
| Inversion | 0.7 | 0.7 | 0.7 | 0.7 | 0.7 |
| Duplication | -1 | -1 | -1 | -1 | -1 |
| Intra-fusion | 0.05 | 0.3* | 0.05 | 0.05 | 0.3* |
| Inter-translocation | 0.05 | 0.65* | 0.05 | 0.05 | 0.65* |
| Control-group | 0% | 0% | 1% | 1% | 1% |
| SV masking | all | all | all | all | all |
| Self-molecules count | 5 | 5 | 5 | 5 | 5 |
| SV chimeric score | show not failing | show not failing | show not failing | show not failing | show not failing |
| Copy Number Variants | 0.99 | 0.99 | 0.99 | 0.99 | 0.99 |
| Copy Number Variants masking | non-masked only | non-masked only | non-masked only | all | all |
| Aneuploidy | 0.95 | 0.95 | 0.95 | 0.95 | 0.95 |

The settings of the several filters are shown in this table. The following positive predictive values are recommended in Bionano Access v.1.7: insertion/deletion = 0, inversion = 0.7, duplications = -1, intra-fusion = 0.05, inter-translocation = 0.05, CNV = 0.99, aneuploidy = 0.95. Recommended Bionano Access v.1.6 settings are the same except: intra-fusion = 0.3 and inter-translocation = 0.65. A control database was used for filtering. Variants with an occurrence of 0 or 1% in the database were considered for analysis. CNV-sensitive regions were masked with filter settings A, B and C. Other settings that were applied to all were: SV-masking = all, self-molecules-counts = 5, SC-chimeric-score-show-not-failing.

#### Supplemental Table 2: number of aberrations with the several filter-sets, *de-novo* pipeline

| Type of aberration | Number of aberrations (Mean/SD) | | Filterset A (Mean/SD) | | Filterset B (Mean/SD) | | Filterset C (Mean/SD) | | Filterset D (Mean/SD) | | Filterset E (Mean/SD) | |
|---|---|---|---|---|---|---|---|---|---|---|---|---|
| **Insertion** | 7090.4 | 2588.4 | 8.3 | 2.9 | 8.3 | 2.9 | 15.2 | 4.3 | 15.2 | 4.3 | 15.2 | 4.3 |
| **Deletion** | 3538.3 | 1547.2 | 12.9 | 3.7 | 12.9 | 3.7 | 25.1 | 4.4 | 25.1 | 4.4 | 25.1 | 4.4 |
| **Inversion** | 207.2 | 24.1 | 0.2 | 0.5 | 0.2 | 0.5 | 0.4 | 0.7 | 0.4 | 0.7 | 0.4 | 0.7 |
| **Duplication** | 119.4 | 53.1 | 2.2 | 1.7 | 2.2 | 1.7 | 2.9 | 2.3 | 2.9 | 2.3 | 2.9 | 2.3 |
| **Intra-Fusion** | 99.0 | 57.0 | 4.7 | 5.7 | 1.0 | 1.2 | 5.7 | 5.7 | 5.7 | 5.7 | 1.2 | 1.4 |
| **Inter-Translocation** | 61.6 | 17.6 | 0.8 | 1.3 | 0.6 | 1.3 | 1.1 | 1.5 | 1.1 | 1.5 | 0.7 | 1.3 |
| **AOH/LOH** | 10.1 | 3.0 | 10.1 | 3.0 | 10.1 | 3.0 | 10.1 | 3.0 | 10.1 | 3.0 | 10.1 | 3.0 |
| **CNV gain** | 28.6 | 9.7 | 2.2 | 2.6 | 2.2 | 2.6 | 2.2 | 2.6 | 2.6 | 2.7 | 2.6 | 2.7 |
| **CNV loss** | 31.7 | 3.9 | 1.1 | 1.9 | 1.1 | 1.9 | 1.1 | 1.9 | 1.2 | 1.9 | 1.2 | 1.9 |
| **aneuploidy Gain** | 0.1 | 0.3 | 0.1 | 0.3 | 0.1 | 0.3 | 0.1 | 0.3 | 0.1 | 0.3 | 0.1 | 0.3 |
| **aneuploidy Loss** | 0.1 | 0.3 | 0.06 | 0.2 | 0.06 | 0.2 | 0.06 | 0.2 | 0.06 | 0.2 | 0.06 | 0.2 |

In the first column, the number of aberrations (mean, SD) after *de-novo* assembly are shown without filtering. In the other columns, the number of aberrations (mean, SD) are depicted for each filter set. The different settings are presented in table 1.

#### Supplemental Table 3: number of aberrations with the several filter-sets, RVA

| Type of aberration | Number of aberrations (Mean/SD) | | Filterset A (Mean/SD) | | Filterset B (Mean/SD) | | Filterset C (Mean/SD) | | Filterset D (Mean/SD) | | Filterset E (Mean/SD) | |
|---|---|---|---|---|---|---|---|---|---|---|---|---|
| **Insertion** | 771.2 | 64.0 | 6.6 | 3.3 | 6.6 | 3.3 | 10.4 | 2.9 | 10.4 | 2.9 | 10.4 | 2.9 |
| **Deletion** | 751.5 | 34.2 | 14.1 | 5.3 | 14.1 | 5.3 | 21.4 | 5.4 | 21.4 | 5.4 | 21.4 | 5.4 |
| **Inversion** | 198.5 | 38.8 | 0.0 | 0.0 | 0.0 | 0.0 | 0.0 | 0.0 | 0.0 | 0.0 | 0.0 | 0.0 |
| **Duplication** | 237.0 | 99.8 | 2.1 | 1.1 | 2.1 | 1.1 | 3.6 | 1.8 | 3.6 | 1.8 | 3.6 | 1.8 |
| **Intra-Fusion** | 108.2 | 50.3 | 2.6 | 1.2 | 0.6 | 0.8 | 3.9 | 1.8 | 3.9 | 1.8 | 0.6 | 0.8 |
| **Inter-Translocation** | 58.9 | 27.7 | 0.5 | 1.4 | 0.1 | 0.3 | 0.6 | 1.4 | 0.6 | 1.4 | 0.1 | 0.3 |
| **CNV gain** | 25.3 | 8.5 | 1.5 | 1.8 | 1.5 | 1.8 | 1.5 | 1.8 | 1.6 | 2.1 | 1.6 | 2.1 |
| **CNV loss** | 37.8 | 12.6 | 1.1 | 2.3 | 1.1 | 2.3 | 1.1 | 2.3 | 1.1 | 2.3 | 1.1 | 2.3 |
| **aneuploidy Gain** | 0.2 | 0.4 | 0.2 | 0.4 | 0.2 | 0.4 | 0.2 | 0.4 | 0.2 | 0.4 | 0.2 | 0.4 |
| **aneuploidy Loss** | 0.2 | 0.4 | 0.1 | 0.3 | 0.1 | 0.3 | 0.1 | 0.3 | 0.1 | 0.3 | 0.1 | 0.3 |

In the first column, the number of aberrations (mean, SD) after rare variant analysis are shown without filtering. In the other columns, the number of aberrations (mean, SD) are depicted for each filter set. The different settings are presented in table 1.

#### Supplemental Information 2: BED files

#### Myeloid BED file

| Myeloid BED file GRCh37 (hg19) | | | |
|---|---|---|---|
| ***Gene*** | **Chr** | **Chr Start** | **Chr End** |
| *PRDM16* | 1 | 2985741 | 3355185 |
| *CSF3R* | 1 | 36931643 | 36948915 |
| *MPL* | 1 | 43803474 | 43820135 |
| *RBM15* | 1 | 110881944 | 110889303 |
| *NRAS* | 1 | 115247084 | 115259515 |
| *DNMT3A* | 2 | 25455829 | 25565459 |
| *SF3B1* | 2 | 198256697 | 198299771 |
| *IDH1* | 2 | 209100952 | 209119806 |
| *RASSF1* | 3 | 50367216 | 50378367 |
| *GATA2* | 3 | 128198264 | 128212030 |
| *MECOM* | 3 | 168801286 | 169381563 |
| *FIP1L1* | 4 | 54243819 | 54326103 |
| *CHIC2* | 4 | 54875957 | 54930815 |
| *PDGFRA* | 4 | 55095263 | 55164412 |
| *KIT* | 4 | 55524094 | 55606881 |
| *TET2* | 4 | 106067841 | 106200960 |
| *PDGFRB* | 5 | 149493401 | 149535422 |
| *RPS14* | 5 | 149823791 | 149829319 |
| *RANBP17* | 5 | 170288895 | 170727019 |
| *NPM1* | 5 | 170814707 | 170837888 |
| *DEK* | 6 | 18224399 | 18264799 |
| *CUX1* | 7 | 101459183 | 101927250 |
| *EZH2* | 7 | 148504463 | 148581441 |
| *MNX1* | 7 | 156786744 | 156802129 |
| *MNX1* | 7 | 156797546 | 156803347 |
| *PCM1* | 8 | 17780365 | 17887457 |
| *FGFR1* | 8 | 38268655 | 38326352 |
| *KAT6A* | 8 | 41786996 | 41909505 |
| *RUNX1T1* | 8 | 92967194 | 93115454 |
| *JAK2* | 9 | 4985244 | 5128183 |
| *MLLT3* | 9 | 20341662 | 20622514 |
| *ABL1* | 9 | 133589267 | 133763062 |
| *NUP214* | 9 | 134000980 | 134110057 |
| *MLLT10* | 10 | 21823573 | 22032559 |
| *HRAS* | 11 | 532241 | 535550 |
| *NUP98* | 11 | 3696239 | 3819022 |
| *WT1* | 11 | 32409321 | 32457081 |
| *CCND1* | 11 | 69455872 | 69469242 |
| *PICALM* | 11 | 85668213 | 85780139 |
| *ZBTB16* | 11 | 113931287 | 114121397 |
| *KMT2A* | 11 | 118307204 | 118397539 |

| | | | |
|---|---|---|---|
| *CBL* | 11 | 119076985 | 119178859 |
| *ETV6* | 12 | 11802787 | 12048325 |
| *ETNK1* | 12 | 22778075 | 22843608 |
| *KRAS* | 12 | 25357722 | 25403854 |
| *PTPN11* | 12 | 112856535 | 112947717 |
| *NCOR2* | 12 | 124808956 | 125052010 |
| *FLT3* | 13 | 28577410 | 28674729 |
| *RB1* | 13 | 48877882 | 49056026 |
| *PML* | 15 | 74287013 | 74340155 |
| *IDH2* | 15 | 90627211 | 90645708 |
| *CREBBP* | 16 | 3775055 | 3930121 |
| *MYH11* | 16 | 15796991 | 15950887 |
| *FUS* | 16 | 31191430 | 31206192 |
| *CBFB* | 16 | 67063049 | 67134958 |
| *PRPF8* | 17 | 1553922 | 1588176 |
| *TP53* | 17 | 7571719 | 7590868 |
| *NF1* | 17 | 29421944 | 29704695 |
| *RARA* | 17 | 38465422 | 38513895 |
| *SRSF2* | 17 | 74730196 | 74733493 |
| *SETBP1* | 18 | 42260137 | 42648475 |
| *CALR* | 19 | 13049413 | 13055304 |
| *CEBPA* | 19 | 33790839 | 33793470 |
| *ASXL1* | 20 | 30946146 | 31027122 |
| *RUNX1* | 21 | 36160097 | 36421595 |
| *ERG* | 21 | 39739182 | 40033704 |
| *U2AF1* | 21 | 44513065 | 44527688 |
| *BCR* | 22 | 23522551 | 23660224 |
| *MKL1* | 22 | 40806291 | 41032690 |
| *ZRSR2* | 23 | 15808573 | 15841382 |
| *BCOR* | 23 | 39910498 | 40036582 |
| *STAG2* | 23 | 123094409 | 123236505 |

#### Mature B cell neoplasm BED file

| **Mature B cell neoplasm BED file GRCh37 (hg19)** | | | |
|---|---|---|---|
| ***Gene*** | **Chr** | **Chr Start** | **Chr End** |
| *FAF1* | 1 | 50906935 | 51425936 |
| *CDKN2C* | 1 | 51434366 | 51440309 |
| *BCL10* | 1 | 85731459 | 85742587 |
| *MTF2* | 1 | 93544792 | 93604638 |
| *TMED5* | 1 | 93615299 | 93646246 |
| *FAM46C* | 1 | 118148604 | 118171011 |
| *CKS1B* | 1 | 154947118 | 154951725 |
| *ACP1* | 2 | 264868 | 278282 |
| *MYCN* | 2 | 16080559 | 16087129 |
| *ALK* | 2 | 29415639 | 30144477 |

| | | | |
|---|---|---|---|
| *MSH2* | 2 | 47630205 | 47710367 |
| *BCL11A* | 2 | 60684328 | 60780633 |
| *REL* | 2 | 61108629 | 61155291 |
| *XPO1* | 2 | 61705068 | 61765418 |
| *CXCR4* | 2 | 136871918 | 136875725 |
| *SF3B1* | 2 | 198256697 | 198299771 |
| *MYD88* | 3 | 38179969 | 38184510 |
| *SETD2* | 3 | 47057897 | 47205467 |
| *CDC25A* | 3 | 48198667 | 48229801 |
| *ATRIP* | 3 | 48488113 | 48507708 |
| *FOXP1* | 3 | 71003864 | 71633140 |
| *PIK3A-PIK3CA* | 3 | 178866310 | 178952497 |
| *BCL6* | 3 | 187439164 | 187463513 |
| *TP63* | 3 | 189349215 | 189615068 |
| *FGFR3* | 4 | 1795039 | 1810599 |
| *WHSC1* | 4 | 1873123 | 1983934 |
| *NPM1* | 5 | 170814707 | 170837888 |
| *DUSP22* | 6 | 292056 | 351355 |
| *IRF4* | 6 | 391738 | 411443 |
| *CCND3* | 6 | 41902671 | 42016610 |
| *SEC63* | 6 | 108188959 | 108279482 |
| *FOXO3* | 6 | 108881025 | 109005971 |
| *MYB* | 6 | 135502452 | 135540311 |
| *POT1* | 7 | 124462439 | 124570037 |
| *BRAF* | 7 | 140433812 | 140624564 |
| *TRAIL-TNFRSF10A* | 8 | 23048969 | 23082680 |
| *TRIM35* | 8 | 27142403 | 27168834 |
| *MYC* | 8 | 128748314 | 128753680 |
| *PVT1* | 8 | 128806778 | 129113499 |
| *MAFA* | 8 | 144510229 | 144512602 |
| *NOTCH1* | 9 | 139388884 | 139440238 |
| *PTEN* | 10 | 89623194 | 89731687 |
| *PIK3A* | 10 | 98353068 | 98480279 |
| *CCND1* | 11 | 69455872 | 69469242 |
| *MRE11A* | 11 | 94150468 | 94227040 |
| *BIRC3* | 11 | 102188180 | 102210135 |
| *ATM* | 11 | 108093558 | 108239826 |
| *H2AFX* | 11 | 118964584 | 118966177 |
| *CCND2* | 12 | 4382901 | 4414522 |
| *RB1* | 13 | 48877882 | 49056026 |
| *DLEUregion* | 13 | 50456688 | 51417885 |
| *DLEU2* | 13 | 50556687 | 50699677 |
| *BCMS-DLEU* | 13 | 50975361 | 51205735 |
| *TGDS* | 13 | 95226307 | 95248529 |
| *MGA* | 15 | 41952609 | 42062141 |
| *IGH* | 16 | 31973408 | 31973499 |
| *MAF* | 16 | 79627744 | 79634622 |

| | | | |
|---|---|---|---|
| *PLCG2* | 16 | 81812898 | 81991899 |
| *TP53* | 17 | 7571719 | 7590868 |
| *CLTC* | 17 | 57697049 | 57774317 |
| *MALT1* | 18 | 56338617 | 56417371 |
| *BCL2* | 18 | 60790578 | 60986613 |
| *MAFB* | 20 | 39314516 | 39317876 |
| *PRAME* | 22 | 22890117 | 22901768 |
| *BTK* | 23 | 100604434 | 100641212 |

#### ALL BED file

| **ALL BED file GRCh37 (hg19)** | | | |
|---|---|---|---|
| ***Gene*** | **Chr** | **Chr Start** | **Chr End** |
| *TAL1* | 1 | 47681961 | 47698007 |
| *STIL* | 1 | 47715810 | 47779819 |
| *JAK1* | 1 | 65298905 | 65432187 |
| *MEF2D* | 1 | 156433512 | 156460391 |
| *PBX1* | 1 | 164528596 | 164821060 |
| *ABL2* | 1 | 179068461 | 179112224 |
| *FHIT* | 3 | 59735035 | 61237133 |
| *PDGFRA* | 4 | 55095263 | 55164412 |
| *AFF1* | 4 | 87856153 | 88062206 |
| *DUX4* | 4 | 190998875 | 191000255 |
| *IL3* | 5 | 131396346 | 131398896 |
| *CSF1R* | 5 | 149432853 | 149492935 |
| *PDGFRB* | 5 | 149493401 | 149535422 |
| *EBF1* | 5 | 158122922 | 158526788 |
| *TLX3* | 5 | 170736287 | 170739138 |
| *RUNX2* | 6 | 45296053 | 45518819 |
| *MYB* | 6 | 135502452 | 135540311 |
| *MLLT4-AS1* | 6 | 168224569 | 168227476 |
| *MLLT4* | 6 | 168227670 | 168372700 |
| *HOXA13* | 7 | 27236498 | 27239725 |
| *IKZF1* | 7 | 50344377 | 50472798 |
| *TRB* | 7 | 142197571 | 142198055 |
| *MYC* | 8 | 128748314 | 128753680 |
| *JAK2* | 9 | 4985244 | 5128183 |
| *MLLT3* | 9 | 20344967 | 20622514 |
| *CDKN2A* | 9 | 21967750 | 21994490 |
| *CDKN2B* | 9 | 22002901 | 22009312 |
| *PAX5* | 9 | 36833271 | 37034476 |
| *ABL1* | 9 | 133589267 | 133763062 |
| *NUP214* | 9 | 134000980 | 134109091 |
| *MLLT10* | 10 | 21823573 | 22032559 |
| *PTEN* | 10 | 89623194 | 89731687 |

| | | | |
|---|---|---|---|
| *BLNK* | 10 | 97951454 | 98031333 |
| *TLX1* | 10 | 102891060 | 102897546 |
| *ADD3* | 10 | 111756107 | 111895323 |
| *NUP98* | 11 | 3696239 | 3819022 |
| *LMO1* | 11 | 8245850 | 8290182 |
| *LMO2* | 11 | 33880122 | 33913836 |
| *RAG1* | 11 | 36589563 | 36601310 |
| *RAG2* | 11 | 36613492 | 36619829 |
| *KMT2A* | 11 | 118307204 | 118397539 |
| *ZNF384* | 12 | 6775642 | 6798738 |
| *ETV6* | 12 | 11802787 | 12048325 |
| *BTG1* | 12 | 92534053 | 92539673 |
| *RB1* | 13 | 48877882 | 49056026 |
| *TRA* | 14 | 22180545 | 22181065 |
| *TRD* | 14 | 22564301 | 22564896 |
| *BCL11B* | 14 | 99635624 | 99737822 |
| *PAR1* | 15 | 25380788 | 25383200 |
| *IGH* | 16 | 31973408 | 31973499 |
| *NF1* | 17 | 29421944 | 29704695 |
| *IKZF3* | 17 | 37913967 | 38020441 |
| *HLF* | 17 | 53342320 | 53402426 |
| *TCF4* | 18 | 52889562 | 53303224 |
| *TCF3* | 19 | 1609291 | 1652326 |
| *MLLT1* | 19 | 6210391 | 6279959 |
| *EPOR* | 19 | 11487880 | 11495018 |
| *JAK3* | 19 | 17935592 | 17958841 |
| *ELL* | 19 | 18553472 | 18632937 |
| *RUNX1* | 21 | 36160097 | 36421595 |
| *ERG* | 21 | 39739182 | 40033704 |
| *TMPRSS2* | 21 | 42836477 | 42880085 |
| *BCR* | 22 | 23522551 | 23660224 |
| *CRLF2* | 23 | 1314886 | 1331616 |
| *CSF2RA* | 23 | 1387692 | 1428828 |
| *IL3RA* | 23 | 1455508 | 1501582 |
| *P2RY8* | 23 | 1581465 | 1656037 |

**Supplemental Information 3: OGM procedure, OGM-detected aberrations and confirmation of these aberrations with the SOC methods.**

After collection of the samples, 15 µl stabilizing buffer (Bionano) was added to 1 ml of bone marrow aspirate (BMA). Samples were stored at -80°C until transported on dry ice to a Bionano certified service provider lab (INRAe, Clermont-Ferrand, France), where the OGM procedure was performed following the manufacturer’s instructions. For each sample, 1 mL of BMA was used to purify ultra-high molecular weight (UHMW) DNA using the Bionano Prep SP BMA DNA Isolation kit. Molecules were labeled with the DLS (Direct Label and Stain) DNA Labeling Kit. Saphyr chips were run to reach a minimum yield of 1300 Gb. The *de novo* assembly, RVA and variant annotation pipeline were executed on Solve v.3.7 (Bionano). The *de novo* assembly was run with a coverage of ~200x and the RVA with ~400x mapped reads. Reporting and direct visualization of structural variants was done on Access v.1.7 (Bionano). Filtering of the structural variants was optimized by testing several filter settings. OGM was successfully executed for all 18 bone marrow samples, and 23 aberrations were detected in the regions of interest. In total, 13/23 OGM findings could be confirmed with the SOC methods, and no SOC-detected aberrations were missed by OGM (supplemental table 4).

### Supplemental Table 4: OGM-detected aberrations

| ID | Referral reason | OGM (splitted per aberration) | *De novo* / RVA / Both | Reason not detected | SOC | Identified with SOC |
|---|---|---|---|---|---|---|
| BMA1 | AML/MDS/MPN | ogm[GRCh37]1q21.1q21.1(144452084_249237532)x2~3 | both | | Karyotyping | 46,XY,der(13)t(1;13)(q12;p11) |
| | | ogm[GRCh37]17p13.1(7,545,377_7,588,071)x1 | *de novo* | RVA only aberrations >5kb | Could not be confirmed | - |
| BMA2 | CML | ogm[GRCh37]t(9;22)(q34;q11.2)(133619362_23603312) | both | | Karyotyping | 46,XY,t(9;22)(q34;q11.2) |
| BMA3 | AML/MDS | ogm[GRCh37]17p13.1(7,545,377_7,588,071)x1 | *de novo* | RVA only aberrations >5kb | Could not be confirmed | - |
| BMA4 | MDS/MPN | ogm[GRCh37](X,1-22)x2 (no aberration) | both | | SNP-arrays | arr(X,1-22)x2 |
| BMA5 | MDS | ogm[GRCh37](X,Y)x1,(1-22)x2 (no aberration) | both | | SNP-arrays | arr(X,Y)x1,(1-22)x2 |
| BMA6 | MDS | ogm[GRCh37]17p13.1(7,545,377_7,588,071)x1 | *de novo* | RVA only aberrations >5kb | Could not be confirmed | - |
| BMA7 | MDS/MF | ogm[GRCh37]inv(3)(q25.33q26.2)(160,014,744_168,882,939) | both | | Could not be confirmed | - |
| | | ogm[GRCh37]3q25.33(159,902,689-160,014,744)x1 | both | | Could not be confirmed | - |
| | | ogm[GRCh37]3q26.2(168,882,939-168,907,480)x1 | both | | Could not be confirmed | - |
| BMA8 | CML | ogm[GRCh37]t(9;22)(q34;q11.2) | both | | Karyotyping | 46,XX,t(9;22)(q34;q11.2) |
| | | ogm[GRCh]22q11.23;q12.1)(23,734,214;29,424,129)x1 | both | | SNP-arrays | arr[GRCh37] 22q11.23q12.1(23632968_29447635)x1 |
| BMA9 | MDS | ogm[GRCh37]chr21x3 | both | | SNP-arrays | arr(21)x3 |
| | | ogm[GRCh37]inv(15)(q24.1q24.1)(72,959,741_74,362,190) | both | | Could not be confirmed | - |
| | | ogm[GRCh37]3p13(71086423_71375386)x1 | RVA | | Could not be confirmed | - |
| BMA10 | MDS/MPN | ogm[GRCh37]9p24.3p13.3(1_35889394)x2 hmz | *de novo* | RVA not able to detect hmz | SNP-arrays | arr[GRCh37] 9p24.3p13.3(1_35889394)x2 hmz, |
| | | ogm[GRCh37]14q11.2q12(20421719_32361050)x1 | both | | SNP-arrays | 14q11.2q12(20421719_32361050)x1 |
| BMA11 | ALL/CLL | ogm[GRCh37](X,Y)x1,(1-22)x2 (no aberration) | both | | Karyotyping | 46,XY |
| BMA12 | (B)-CLL | ogm[GRCh37]chr12x3 | both | | SNP-arrays | arr(12)x3 |
| BMA13 | MDS | ogm[GRCh37] 20q11.22q13.13(34259243_49502193)x1 | both | | SNP-arrays | arr[GRCh37] 20q11.22q13.13(34259243_49502193)x1 |
| BMA14 | ? | ogm[GRCh37](X,Y)x1,(1-22)x2 (no aberration) | both | | Karyotyping | 46,XY |
| BMA15 | MDS/AML | ogm[GRCh37] fus(5;5)(q14.2;q33.3) | RVA | 5% aberrant cells non-cultured | Karyotyping, SNP-arrays | 46,XX,5q-[5]/46,XX[5] (only OGM RVA) |
| BMA16 | HCL (CLL) | ogm[GRCh37](X)x1,(1-22)x2 | both | | Karyotyping | 45,X,-Y[13]/46,XY[7] |
| | | ogm[GRCh37]8p23.3p22(11805_15540773)x1~2 | both | | SNP-arrays | arr[GRCh37] (Y)x0,8p23.3p22(1_15547178)x1, 8q13.2q24.3(69538276_146364022)x3 |
| | | ogm[GRCh37]fus(8;8)(p22;q13.2) | both | | Karyotyping (1:250 metaphases) | fus(8;8)(p22;q13.2),t(14;17)(q32.33;q25.3) |
| | | ogm[GRCh37]t(14;17)(q32.22;q25.3) (106,249,815;80,915,618) | both | Disputable OGM breakpoint | Could not be confirmed | - |
| BMA17 | MDS | OGM[GRCh37]ins(12p13.2)(11,889,007_11,895,594) | *de novo* | RVA only aberrations >5kb | Could not be confirmed | - |
| BMA18 | AML | ogm[GRCh37](X,(1-22)x2 | both | | | 46,XX[20] |

All OGM aberrations detected in a selected group of leukemia patients are depicted in this table. The table also shows if the OGM aberrations are confirmed with the SOC methods.

Abbreviations: ID = sample identification, AML = acute myeloid leukemia, MDS = myelodysplastic syndrome, MPN = myeloproliferative neoplasms, CML = chronic myeloid leukemia, MF = myelofibrosis, ALL = acute lymphoblastic leukemia , CLL = chronic lymphoblastic leukemia, (B)-CLL = B cell acute lymphoblastic leukemia, HCL = hairy cell leukemia, SOC = standard-of-care, OGM = optical genome mapping.

A homozygous region (ogm[GRCh37]9p24.3p13.3(1_35889394)x2 hmz) in sample BMA10 with referral reason myelodysplastic syndrome / myeloproliferative neoplasms was only detected with the *de novo* pipeline because the RVA pipeline cannot detect homozygous regions (Bionano, 30110 Rev K).

A 1.2 kb heterozygous 17p13.1 deletion (ogm[GRCh37]17p13.1(7,545,377_7,588,071)x1) in a 42 kb region that includes *TP53* in samples BMA1, BMA3 and BMA6 and a 1.4 kb heterozygous 12p13.2 insertion in a 6.6 kb region that includes *ETV6* (OGM[GRCh37]ins(12p13.2)(11,889,007_11,895,594)) in sample BMA17 with referral reason myelodysplastic syndrome were only detected with the *de novo* pipeline. The aberrations in BMA1, BMA3, BMA6 and BMA17 were below the resolution of the RVA pipeline (5 kb, Bionano, 30110 Rev K).

A fusion of chromosome 5 (ogm[GRCh37] fus(5;5)(q14.2;q33.3)) in patient BMA15, with referral reason myelodysplastic syndrome / acute myeloid leukemia was only detected with the RVA pipeline. Karyotyping of the aberration identified it as a chromosome 5q deletion in 5 of 10 metaphases. Using SNP-array on non-cultured cells, the estimated percentage of aberrant cells was 5%, which is below the threshold of the *de novo* pipeline (Bionano, 30110 Rev K).

In patient BMA9, with referral reason myelodysplastic syndrome, a 288 kb deletion on chromosome 3 (ogm[GRCh37]3p13(71086423_71375386)x1) with an allele frequency of 3% was only detected with the RVA pipeline. An allele frequency of 3% is below the resolution of the *de novo* pipeline (Bionano, 30110 Rev K). All other aberrations were detected with both pipelines.

In patient BMA16, with referral reason hairy cell leukemia, OGM identified a loss of the p-arm of chromosome 8p23.3p22 (ogm[GRCh37]8p23.3p22(11805_15540773)x1~2) and a chromosome 8 fusion (ogm[GRCh37]fus(8;8)(p22;q13.2)) (table). SNP-array detected a gain and loss of chromosome 8 (arr[GRCh37](Y)x0,8p23.3p22(1_15547178)x1, 8q13.2q24.3(69538276_146364022)x3), and we performed karyotyping to confirm the fusion. The fusion was detected in only 1 of 250 metaphases, probably due to the loss of aberrant mature B cells in the culturing procedure (1). The clinical significance of the fusion is unknown (2,3).

In patient BMA1, with referral reason acute myeloid leukemia / myelodysplastic syndrome / myeloproliferative neoplasms, karyotyping confirmed a gain of 1q21.1q21.1 (ogm[GRCh37]1q21.1q21.1(144452084_249237532)x2~3) as a 46,XY,der(13)t(1;13)(q12;p11) with karyotyping.

**References Supplemental Information 3**

1. Troussard X, Cornet E. Hairy cell leukemia 2018: update on diagnosis, risk-stratification, and treatment. Am J Hematol. 2017;12:1382-1390.

2. The 1000 Genomes Project Consortium. A global reference for human genetic variation. Nature 2015;526(7571):68-74.

3. Edelmann J, Holzmann K, Miller F, Winkler D, Bühler A, Zenz T et al. High-resolution genomic profiling of chronic lymphocytic leukemia reveals new recurrent genomic alterations. Blood 2012;120(24):4783-4794.

#### Supplemental Information 4:

#### Supplemental Table 5: Crispr locations.

| ID | Additional putative OGM aberration | Gene | crRNA GRCh37 hg19 |
|---|---|---|---|
| **BMA1** | OGM[GRCh37]17p13.1(7,545,377_7,588,071)x1 | *TP53* | chr17:7,543,974-7,543,993 + |
| **BMA3** | OGM[GRCh37]17p13.1(7,545,377_7,588,071)x1 | *TP53* | chr17:7,559,625-7,559,644 - |
| **BMA6** | OGM[GRCh37]17p13.1(7,545,377_7,588,071)x1 | *TP53* | chr17:7,556,557-7,556,576 + |
| | | | chr17:7,576,982-7,577,001 - |
| | | | chr17:7,572,406-7,572,425 + |
| | | | chr17:7,589,483-7,589,502 - |
| **BMA9** | OGM[GRCh37]3p13(71,086,423_71,375,386)x1 | *FOXP1* | chr3:71,082,359-71,082,378 + |
| | | | chr3:71,101,049-71,101,068 – |
| | | | chr3:71,082,955-71,082,974 + |
| | | | chr3:71,101,401-71,101,420 – |
| | | | chr3:71,371,938-71,371,957 + |
| | | | chr3:71,384,613-71,384,632 – |
| | | | chr3:71,368,075-71,368,094 + |
| | | | chr3:71,382,344-71,382,363 – |
| **BMA17** | OGM[GRCh37]ins(12p13.2)(11,889,007_11,895,594) | *ETV6* | chr12:11,888,244-11,888,263 + |
| | | | chr12:11,902,197-11,902,216 - |
| | | | chr12:11883363-11883382 + |
| | | | chr12:11910042-11910061 - |
| **BMA16** | OGM[GRCh37]t(14;17)(q32.22;q25.3)(106,249,815;80,915,618) | *IGHG1* | chr17:80967365-80967384 + |
| | | | chr17:80996130-80996149 + |
| | | | chr14:106126211-106126230 - |
| | | | chr14:106100173-106100192 - |
| | | | chr14:106083832-106083851 - |
| **BMA7** | OGM[GRCh37]inv(3)(q25.33;q26.2)(160,014,744_168,882,939) | *MECOM* | chr3:159898486-159898505 + |
| | OGM[GRCh37]3q25.33(159,902,689-160,014,744)x1 | | chr3:160032530-160032549 - |
| | OGM[GRCh37]3q26.2(168,882,939-168,907,480)x1 | | |
| **BMA9** | OGM[GRCh37]inv(15)(q24.1;q24.1)(72,959,741_74,362,190) | *PML* | chr15:72975831-72975850 - |
| | | | chr15:72966403-72966422 - |

For each additional OGM-detected putative aberration, this table lists the sample ID, the genes involved and the GRCh37 coordinates of the crRNA enzymes.
